# Genetically encoded sensors for analysing neurotransmission among synaptically-connected neurons

**DOI:** 10.1101/2022.04.03.486903

**Authors:** Yutaka Shindo, Keita Ashida, Kazuto Masamoto, Hiroyuki Takuwa, Manami Takahashi, Makoto Higuchi, Ryuto Ide, Kohji Hotta, Kotaro Oka

**Author notes:** These authors contributed equally to this work (Yutaka Shindo, Keita Ashida). **Corresponding author** Prof. Kotaro Oka.

## Abstract

Anatomical connectome mapping in small areas of the nervous system as well as large-scale detection of neuronal activity patterns have been respectively achieved; however, it is still challenging to evaluate the functional connections among anatomically-connected neurons in a large-scale nervous system. We have developed a novel method to visualize neurotransmission named Split Protein HEmispheres for REconstitution (Sphere). By splitting a sensor into two fragments and expressing them in pre- and postsynaptic neurons separately, functional neurotransmitter sensors can be reconstituted only at the synapses between those neurons. We developed a Sphere-SF-iGluSnFR to measure glutamate levels, and further demonstrated that this system is functional in cultured cells, worms, and mouse brains. Moreover, this system is applicable to sensors other than glutamate, and colour variants have also been developed. This could allow for brain-wide imaging of functional synaptic transmission among particular neurons and identification of important neuronal circuits in the nervous system.

## Introduction

To understand information processing within the nervous system, it is necessary to understand the connections and transmission among neurons. In recent years, large-scale analysis of the anatomical and functional connections among neurons in the nervous system has been actively pursued. Exhaustive research using electron microscopy^1^ has been able to show the anatomical connections between neurons in parts of the nervous systems of various animals^2–4^. This method has the advantage of obtaining a complete 3D map of synaptic connections within a specific neuronal circuit and visualizing the morphological features of the neurons within it. However, the detected synapses are not always functional, and it is still difficult to analyse large-scale neuronal circuits between multiple regions of the brain and observe synaptic connections *in vivo*. On the other hand, volumetric imaging of neuronal activity using Ca^2+^ imaging has achieved large-scale detection of neuronal activity in zebrafish, worms, and flies^5–9^. The functional connectivity between neurons or among regions of the brain can be estimated by analysing the correlations of their activity patterns, and neurons related to specific behaviours have thus been identified. However, we can only estimate functional connectivity from Ca^2+^ imaging, and it is still uncertain whether the detected functional connections reflect actual synaptic connections unless analysed by other methods such as electron microscopy^10,11^. Therefore, to analyse the functional connections between synapses within a nervous system *in vivo*, a novel neurotransmission imaging technique is necessary.

Neurotransmitter imaging is a key technique to achieve this purpose because neurotransmission at chemical synapses is mediated by neurotransmitters. Recently, various genetically encoded sensors for neurotransmitter imaging have been developed^12–16^. Of these, a glutamate sensor, iGluSnFR, has been used to create mesoscale functional connectome maps^17^. However, because this sensor is expressed not only at synapses, but also throughout the cell membrane, it is difficult to differentiate the anatomical connections among neurons. To visualize synaptic connections between focused neurons *in vivo*, the GFP Reconstitution Across Synaptic Partners (GRASP) system was developed^18,19^. Using enhanced GRASP (eGRASP) technique, anatomical changes in synapses connected to memory engram neurons in the hippocampus have been evaluated^20^. It is now possible to describe changes in the number of synapses and the size of the spines between these neurons that occur with learning. However, only anatomical information is obtained using eGRASP, and the functional connections for neurotransmission are still obscure.

In this study, to link anatomical and functional connections *in vivo*, we developed a novel technique to visualize synaptic neurotransmission. We expected that this could be achieved by combining the properties of neurotransmitter sensors with the GRASP system. We split a genetically encoded neurotransmitter sensor into two fragments so that it would reconstitute only when the fragments encountered each other in a synaptic cleft. By expressing each half of the sensor in separate neurons, the reconstituted sensors are thus localized at the points of attachment of those cells, which are the synapses between pre- and postsynaptic neurons of interest (Fig. 1A). We named this technique Split Protein HEmispheres for REconstitution (Sphere). This technique is expected to enable imaging that integrates anatomical and functional information.

**Figure 1.**
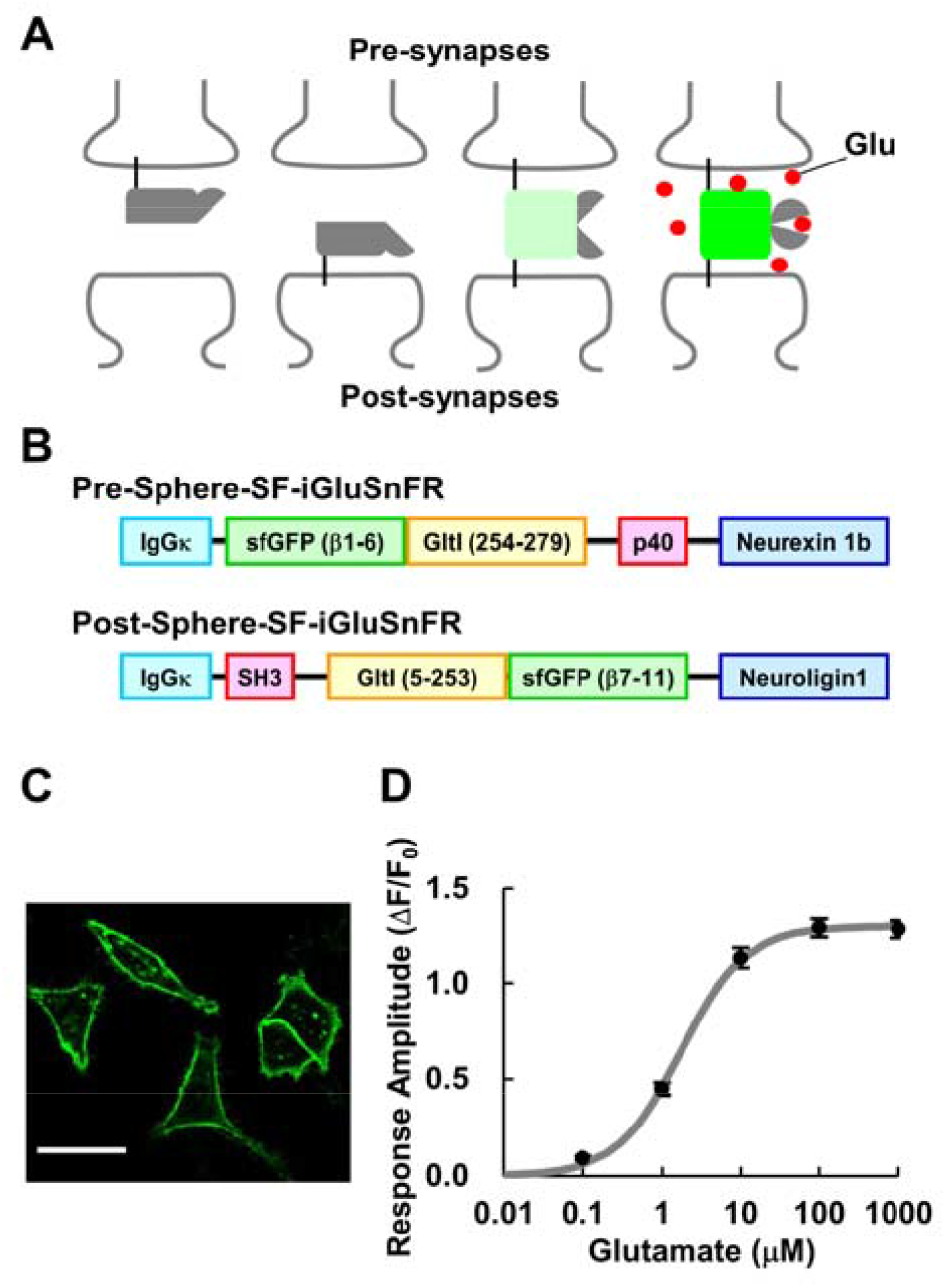
Sensor concept and characterization on a single cell. (A) Concept of the Split Protein HEmispheres for REconstitution (Sphere) technique. The separated sensors were reconstituted and showed fluorescence only at the synapses. (B) Schematic of Pre- and Post-Sphere-SF-iGluSnFR. IgGκ (light blue): IgG kappa secretion tag for extracellular localization; sfGFP (green): fragments of sfGFP, a GFP variant; GltI (orange): fragments of the glutamate sensor domain derived from the *E. Coli* GltI; p40 and SH3 (red): selective binding domain; Neurexin 1b and Neuroligin 1 (blue): the stalk, transmembrane, and intracellular domains of Neurexin 1b and Neuroligin 1 for membrane anchoring. (C) Fluorescence image of Sphere-SF-iGluSnFR reconstituted on the cell membrane of each HeLa cell. Scale bar: 50 μm. (D) Glutamate-dependent response of the Sphere-SF-iGluSnFR (*n* = 104 cells from 8 different experiments). Error bars: standard error of the mean (SEM).

## Results

### Engineering the Sphere technique and its characterization in culture cells

To detect synaptic connections between specific neurons, an effective approach is to separate a fluorescent protein (FP) into two fragments and express each part in pre- and postsynaptic neurons. Each fragment of the FP emits no fluorescence; however, when a complete FP is reconstituted, fluorescence is exhibited only where those neurons attach—a synapse. The GRASP series employs this strategy and has successfully visualized synaptic connections between specific pre- and postsynaptic neurons^18–20^. Applying this strategy to genetically encoded neurotransmitter sensors would enable a visualization of the neurotransmission events between specific pre- and postsynaptic neurons, which is the basis of the Sphere technique developed in this study. A single FP-based glutamate sensor, SF-iGluSnFR^21^, was initially split into two fragments to develop the Sphere-SF-iGluSnFR construct. Based on previous studies, we generated three types of split glutamate sensors: a GRASP-type, a circularly permutated (cp)-type, and a non-circularly permutated (ncp)-type (Supplementary Fig. 1a–c). The GRASP-type of split-GFP construct (Supplementary Fig. 1d) has advantages in minimizing protein interference and aggregation of GFP fragments^22^, and is applicable for multiple FP colour variants^23^. On the other hand, in the case of the glutamate sensors iGluSnFR and G^ncp^-iGluSnFR, EGFP was split at the position between β6 and β7 to maximize the sensor response, and they were arranged in cp or ncp order (Supplementary Fig. 1e and f). These three types of separation were applied to SF-iGluSnFR, and each fragment was replaced with a GFP fragment Pre-and Post-eGRASP to constitute the final Pre- and Post-Sphere-SF-iGluSnFR constructs (Fig. 1B and Supplementary Fig. 1).

Both fragments of the sensor were first expressed together on the surface of HeLa cells to confirm fluorescence and response. The GRASP-type did not reconstitute the fluorescence sensor, but the other two types emitted fluorescence and responded to glutamate concentration changes (Figs. 1C, D and Supplementary Fig. 2). Because binding domains are necessary for the two fragments to be sufficiently close to each other to properly reconstitute the FP^20,24^, mutations in those may affect fluorescence and response of the sensor. Variants of GFP also have significant impact on the sensor response^21^. Therefore, we investigated the effect of these mutations and variants on the sensor response. It was found that the sfGFP-type with Ab1-SH3 and p40 binding domain displayed the best response (Supplementary Fig. 2). The reconstituted Sphere-SF-iGluSnFR showed an almost equivalent response and selectivity to major neurotransmitters compared to the original SF-iGluSnFR (Supplementary Fig. 3).

The reconstitution of the sensors between cells was confirmed using HEK293 cells. Cells expressing Pre-Sphere-SF-iGluSnFR with TagBFP and cells expressing Post-Sphere-SF-iGluSnFR with iRFP were mixed and plated. Green fluorescence from reconstituted Sphere-SF-iGluSnFR proteins was observed only at the attachment sites of those cells (Fig. 2A), and the fluorescence intensity increased in response to glutamate (Fig. 2B). Based on these data, it was concluded that the Sphere technique was successful.

**Figure 2.**
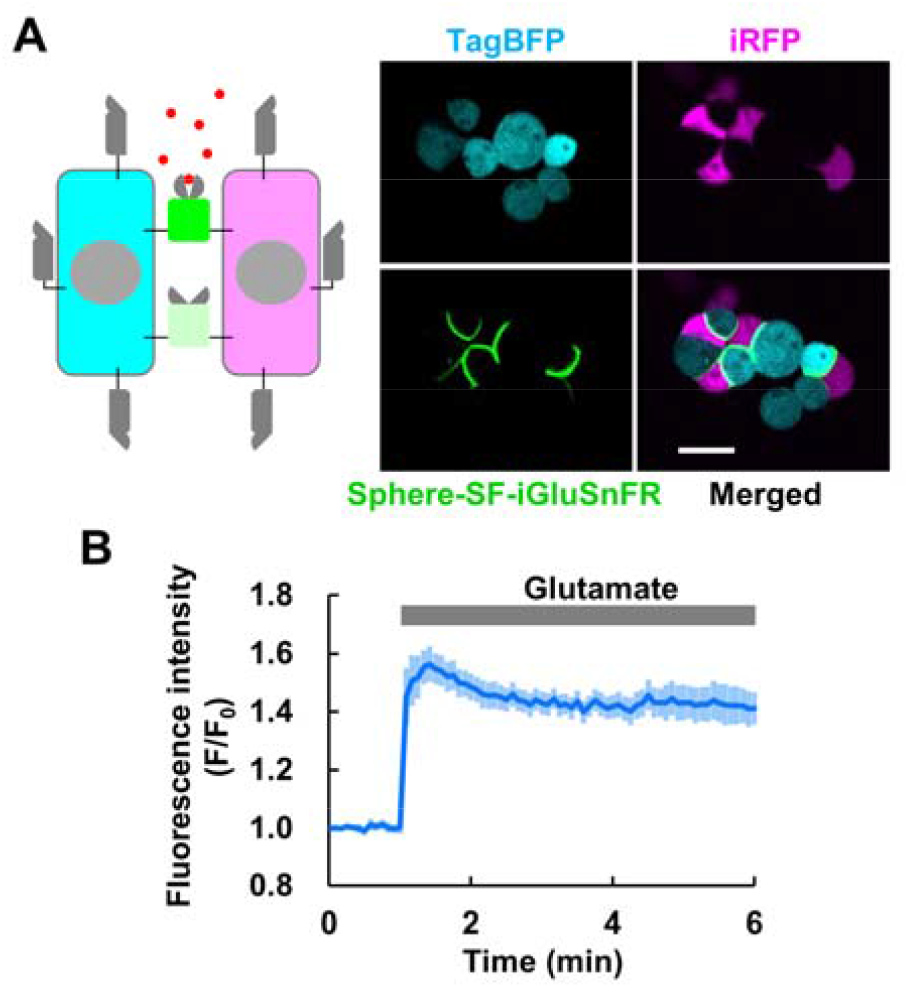
Fluorescence and response of the Sphere-SF-iGluSnFR protein reconstituted between cells. (A) Schematic illustration of the experiment and fluorescence images. TagBFP: a marker for cells expressing Pre-Sphere-SF-iGluSnFR; iRFP: a marker for cells expressing Post-Sphere-SF-iGluSnFR; Sphere-SF-iGluSnFR: the reconstituted Sphere-SF-iGluSnFR; Merged: merged image. Scale bar: 20 μm. (B) Response of Sphere-SF-iGluSnFR reconstituted between cells to 100 μM glutamate (*n* = 16 ROIs from 4 different experiments). Error bars: SEM.

### Application of the Sphere technique to other sensors and other colour variants

To confirm that the Sphere technique is applicable to sensors other than SF-iGluSnFR, it was applied to the genetically encoded GABA (γ-aminobutyric acid) sensor iGABASnFR^16^, and the red colour variant R^ncp^-iGluSnFR1 (Supplementary Fig. 4a and b)^25^. Both sensor fragments were expressed in HeLa cells, and both Sphere-iGABASnFR and Sphere-R^ncp^-iGluSnFR1 successfully emitted fluorescence and responded to GABA and glutamate, respectively, indicating that the complete sensors were reconstituted on the cell membrane (Supplementary Fig. 4c–f). The amplitudes of the observed responses were comparable with those of the original sensors (Supplementary Fig. 4g and h), indicating that the performance of the novel sensors did not deteriorate during fragmentation and reconstitution.

Colour variants of the Sphere-SF-iGluSnFR construct were also developed. By introducing chromophore mutations, cyan and yellow variants were achieved (Supplementary Fig. 5a). These variants emitted fluorescence and properly responded to glutamate (Supplementary Fig. 5b–d). The amplitudes of the responses from the cyan and yellow variants to glutamate were smaller than that of the green variant (Sphere-SF-iGluSnFR); diminished results associated with chromophore mutations was also observed in the original SF-iGluSnFR construct^21^. These results indicate that the Sphere technique can be applied to a broad range of sensors.

### *In vivo* imaging of glutamate transmission between two specific neurons in *Caenorhabditis elegans*

To demonstrate the localization and function of Sphere-SF-iGluSnFR *in vivo*, neurotransmission in *Caenorhabditis elegans* was observed. TagBFP and Pre-SF-iGluSnFR expression was induced in an olfactory sensory neuron (AWC), and mCherry and Post-SF-iGluSnFR were expressed in AIY interneurons. Because the synaptic connections between these neurons have already been anatomically identified^4^ and the odour-dependent decrease in glutamate release from the AWC sensory neuron has been previously demonstrated^26,27^, these neurons are a suitable choice for confirming the performance of Sphere-SF-iGluSnFR. Fluorescence was indeed observed only at the attachment sites of these neurons (Fig. 3A). The AWC neuron is an “off response neuron”, and usually releases glutamate and reduces transmission upon sensing of isoamyl alcohol (IAA)^26,27^. In this experiment, IAA input was done by changing the flow in a microfluidic device^28^. A decrease in the fluorescence of Sphere-SF-iGluSnFR was observed during IAA exposure (Fig. 3B), as was observed with iGluSnFR in previous studies^26,27^. These data clearly indicate that Sphere-SF-iGluSnFR successfully reconstituted glutamate sensor function at the synapses between two specific neurons. Therefore, the functionality of our probe was confirmed *in vivo*.

**Figure 3.**
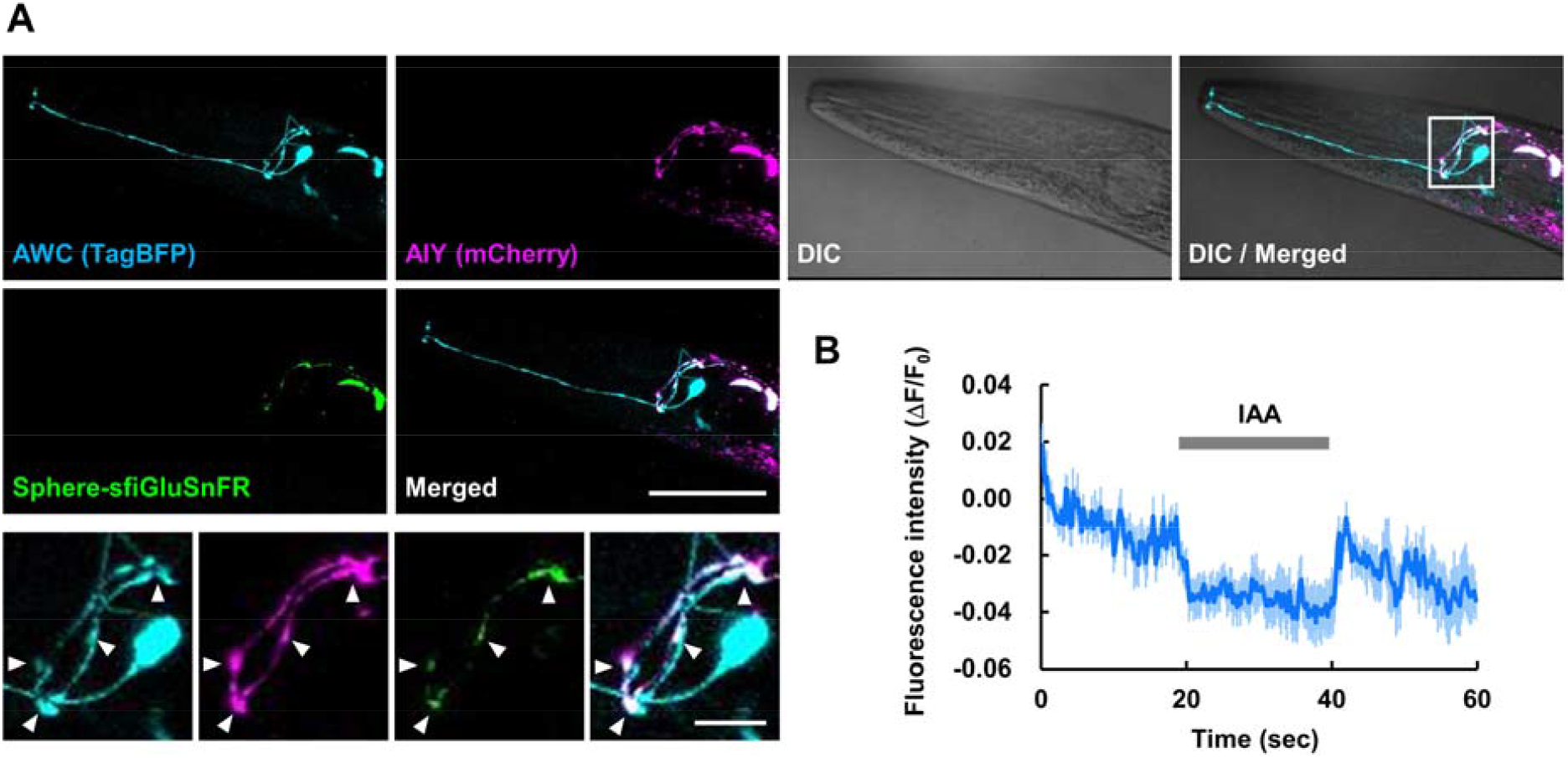
Imaging of glutamate transmission between two specific neurons in *Caenorhabditis elegans*. (A) Fluorescence image of TagBFP as an AWC sensory neuron marker; mCherry as an AIY interneuron marker; and the reconstituted Sphere-SF-iGluSnFR, merged image, and merged image with differential interference contrast (DIC) images obtained via confocal microscopy (upper six panels). Magnified images of the position in the DIC-merged image are included. Arrow heads indicate the positions where fluorescence of the reconstituted Sphere-SF-iGluSnFR was observed (lower four panels). Scale bars: 50 μm for upper panels, 10 μm for lower panels. (B) The average glutamate transmission between an AWC sensory neuron and an AIY interneuron in response to an application of 100 μM isoamyl alcohol (IAA) to *C. elegans*, measured using fluorescence intensity as a result of Sphere-SF-iGluSnFR expression (n = 6 worms). Error bars: SEM.

### *In vivo* imaging of glutamate transmission in mice

The Sphere-SF-iGluSnFR construct was further applied to *in vivo* mouse brain imaging. An adeno-associated virus (AAV) vector encoding Pre- or Post-Sphere-SF-iGluSnFR with marker FP was injected near areas within the cortex (Fig. 4A). Three weeks after injection, two-photon imaging was performed *in vivo*. In the cortex, local interneurons are connected to each other, and ideally Sphere-SF-iGluSnFR expression would localize to the synapses among them. Respective TagBFP and mCherry expression was detected near the injected area, and synapse-like granules of green fluorescence were detected in both areas (Fig. 4B). Spontaneous fluorescent blinking of Sphere-SF-iGluSnFR, which indicates glutamate transmission, was observed through time-lapse imaging (Fig. 4C), and was reduced when the mouse was under anaesthetised conditions (Fig. 4D). Previous studies showed a reduction in neural activity by measuring field potential and electroencephalogram (EEG) imaging^29–31^, which reflects the activity of both local interneurons and projection neurons. On the other hand, our results focus on neurotransmission only among local interneurons and show that they were suppressed under anaesthetised conditions. To compare the glutamatergic synaptic activity in detail, 75 regions of interest (ROIs) that displayed activity while the mouse was awake were selected, and the responses under awake and anaesthetised conditions were compared. The ROIs were clustered based on the similarity of the detected responses, and the responses were plotted in raster plots (Fig. 4E). Multiple synapses were observed to synchronously respond in the same frame under awake conditions, however this was not observed under anaesthetised conditions (red dots in Fig. 4E). The distribution of the number of responding synapses within the same frame also differed under each condition (Fig. 4F). Furthermore, the response frequency decreased in many synapses under anaesthetised conditions (Fig. 4G). Similar results were also observed in another mouse that was investigated in the reverse order, first under anaesthetised conditions and then awake (Supplementary Fig. 6). Relationships between the response patterns and distances between the synapses were calculated for all combinations of the 75 ROIs (Fig. 4H). Some synapses at close distances showed high response similarity, but no consistent response and location trends were observed.

**Figure 4.**
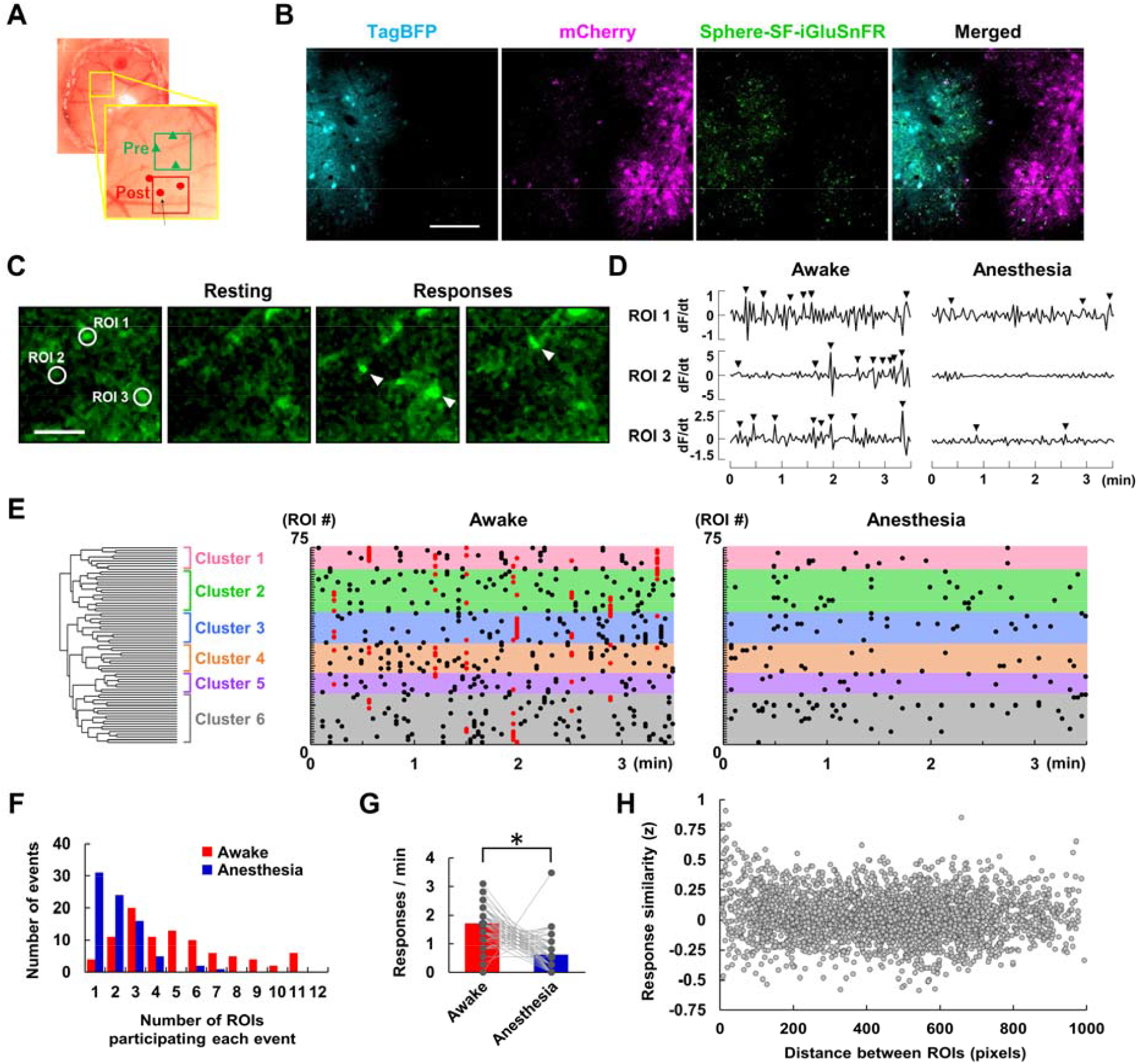
Application of the Sphere-SF-iGluSnFR technique to *in vivo* imaging of glutamate transmission between local interneurons in the cerebral cortex of a mouse brain. (A) Positions of adeno-associated virus (AAV) injection in the mouse brain. (B) Fluorescence images of TagBFP (Pre- marker), mCherry (Post- marker), the reconstituted Sphere-SF-iGluSnFR protein, and the merged image. Scale bar: 100 μm. (C) Representative fluorescence images of Sphere-SF-iGluSnFR responses. Positions of regions of interest (ROIs; left) and images at time frames when some responses were observed. Arrowheads indicate responding points. The responses were observed under awake and anaesthetised conditions at the same position of the brain in the same mouse, and were analysed using the same ROIs. Scale bar: 10 μm. (D) Time-courses of the differential fluorescence intensity (dF/dt) at each ROI. Arrowheads indicate responses detected by our program. (E) Clustering (left) and raster plots of the responses of the Sphere-SF-iGluSnFR at 75 ROIs placed in single time series for 3.5 min (middle and right). The responses were sorted based on the similarity of the responses and were clustered into six groups, which are indicated in different colours. Synchronized responses (more than 10 ROIs responding at the same time point) are indicated by red dots in the raster plots. (F) Histogram of the number of ROIs that responded at the same time frame under awake (red) and anaesthetised (blue) conditions. (G) The number of responses under awake and anesthetised conditions. The grey dots indicate each ROI, and the dots for the same ROI under each condition are connected with a grey line. The red and blue bars indicate the average value of the 75 ROIs under each condition. * indicates *P* > 0.05, Student’s *t*-test. (H) Relationship between the responses and distance between two ROIs. The values were calculated for all combinations of the 75 ROIs, and the response similarity was shown using Fisher Z-transformed values.

## Discussion

In this study, we developed a technique that involves a single FP-based genetically encoded sensor split into two fragments that reconstitutes into a complete sensor at the point of contact between cells. This technique, Sphere, enables the detection of a signal between specific neurons that each express a fragment of the Sphere sensor. This allows for measurement of the neurotransmission between specific pre- and postsynaptic neurons. It was confirmed that Sphere-SF-iGluSnFR localized to the attachment sites between pre- (AWC) and post- (AIY) synaptic neurons and responded to glutamate transmission in *C. elegans*. Although AIY neurons synaptically connect to a number of neurons, the glutamate transmission between specific AWC and AIY neurons could be visualized using Sphere-SF-iGluSnFR because both Pre- and Post- sensor fragments encountered each other and were reconstituted only at the synapses between these neurons. Moreover, Sphere-SF-iGluSnFR was applied to *in vivo* imaging of mouse brains to visualize anaesthesia-induced changes in glutamate transmission specifically among local interneurons. Although recent technological advancements have enabled the analysis of neuronal connections^2,3^ and activities^5,9^, large-scale analysis of information processing among synaptically-connected neuronal circuits *in vivo* has still been a challenge. The Sphere technique developed here would be a powerful tool to visualize neurotransmission between specific neurons within a nervous system. Combining the use of this technique with activity-dependent expression under immediate early gene promoters would help to localize the Sphere sensor to particular synapses related to specific situations^20,32^. Additionally, ratiometric use with another FP might improve the signal-to-noise ratio, allowing for the detection of small signals at synapses during high-speed imaging. Further applications of the Sphere sensors would thus facilitate neuronal imaging.

In addition, we confirmed the functionality of the Sphere-iGABASnFR, which suggests that the Sphere technique is applicable to a number of single-wavelength sensors based on cp-type FPs, such as iATPSnFR for ATP^33^, iGlucoSnFR for glucose^34^, eLACCO1 for lactate^35^. However, it is difficult to apply the Sphere technique to G-protein coupled receptor (GPCR)-based sensors^13–15,36^, due to the fact that Sphere functions by reconstituting an FP in the extracellular space between cells, and FPs in those sensors are embedded in an intracellular GPCR loop.

Cyan, yellow, and red colour variants of the Sphere-SF-iGluSnFR construct were also developed. This enables the combined use of the Sphere probes alongside green fluorescence probes, such as GCaMP^37,38^ to infer the relationship between neurotransmission and neural activity in specific neuronal circuits. The combined use of two colour-variants of Sphere-SF-iGluSnFR might allow for comparing the inputs from two neurons to a single particular neuron. In a previous study, eGRASP was used to label synapses formed between specific pre- and postsynaptic neurons with two colours to compare the morphological changes of synapses among memory engram neurons^20^. The transmission among those neurons could be compared directly using Sphere-SF-iGluSnFR. Further improvements in Sphere probes might enable the simultaneous measurement of excitatory and inhibitory inputs in specific neuronal circuits, such as the central pattern generator (CPG) circuit *in vivo*^39^. We expect that this Sphere technology will substantially advance *in vivo* cellular communications imaging.

## Methods

### Ethics statement

The use of animals and experimental protocols were approved by the Animal Ethics Committee at the University of Electro-Communications (Permit number: A34). All methods were carried out in accordance with the relevant guidelines and regulations. All experimental procedures were conducted according to the Animal Research: Reporting of *In Vivo* Experiments (ARRIVE) guidelines^40^.

### Sensor engineering

The pAAV-CWB-cyan-pre-eGRASP(p32) and pAAV-EWB-DIO-myriRFP670V5-P2A-post-eGRASP vectors were gifts from Dr Bong-Kiun Kaang (Addgene plasmid #111579 and #111585, respectively). The pAAV.CAG.SF-Venus-iGluSnFR.A184V and pAAV.hSynap.iGABASnFR vectors were gifts from Dr Loren Looger (Addgene plasmid #106202 and #112159, respectively). The pDisplay-Gncp-iGluSnFR and pDisplay-R-iGluSnFR1 vectors were gifts from Dr Robert Campbell (Addgene plasmid #107337 and #107335, respectively).

Cyan pre-eGRASP(p32) and myriRFP670V5-P2A-post-eGRASP were inserted in the pcDNA3.1(+) vector (Thermo Fisher Scientific, Waltham, MA, USA) at the NheI-EcoRI and HindIII-XbaI sites, respectively. EGFP (β1–6) with GltI (254–279) and EGFP (β7–11) with GltI (5–253) were amplified from the G^ncp^-iGluSnFR construct by PCR, and were replaced with the FP fragment in Cyan pre-eGRASP(p32) and post-eGRASP via Infusion (Takara Bio, Shiga, Japan) to yield the Pre-Sphere-iGluSnFR(p32) and Post-Sphere-iGluSnFR constructs. To evaluate GFP variants, EGFP fragments from Pre-Sphere-iGluSnFR(p32) and Post-Sphere-iGluSnFR were replaced with GFP from eGRASP, sfGFP, or sfGFP(S72A). To prepare the cp-type and ncp-type shown in Supplementary Fig. 1, the sfGFP (β1–6) with GltI (254–279) and sfGFP (β7–11) with GltI (5–253) in the Pre- and Post-Sphere-SF-iGluSnFR constructs were exchanged with each other. To yield the p40 version of Pre-Sphere-SF-iGluSnFR, p32 in Pre-Sphere-SF-iGluSnFR was mutated via PCR. To prepare the GRASP-type shown in Supplementary Fig. 1a, the FP fragment in Cyan pre-eGRASP was replaced by G^ncp^-iGluSnFR lacking the β11 of EGFP, and the FP fragment in post-eGRASP was replaced with the β11 of EGFP. As markers for transfection, TagBFP or iRFP were bound to the N-terminus of the Pre- and Post-Sphere-SF-iGluSnFR constructs via a P2A linker.

### Fluorescence imaging on cultured cells

The properties of each candidate were evaluated by expressing both sensor fragments on the surface of HeLa cells. HeLa cells were cultured in Dulbecco’s Modified Eagle Medium (DMEM; Thermo Fisher Scientific) supplemented with 10% (v/v) FBS, 50 U/mL penicillin, and 50 μg/mL streptomycin. To evaluate sensor properties, the plasmids encoding Pre- and Post-Sphere sensors were transfected into the HeLa cells by electroporation using a Neon Transfection System (Thermo Fisher Scientific), and the cells were subsequently plated onto a glass bottom dish (Iwaki, Tokyo, Japan). 1–2 days after transfection, the cells were rinsed with Hank’s Balanced Salt Solution (HBSS; Thermo Fisher Scientific) with 10 mM HEPES and a pH adjusted to 7.4. Cells were observed on a FluoView FV1000 confocal laser scanning microscope (Olympus, Tokyo, Japan) equipped with a 60× oil immersion objective (UPlanSApo, NA: 1.35, Olympus). The sensor candidate was excited at 488 nm using an Ar laser (Olympus) through a DM405/488 dichroic mirror (Olympus), and the fluorescence at 500–600 nm through the spectroscopy unit was observed using a photomultiplier (Olympus). Images were obtained every 5 s.

Reconstitution of the Sphere-SF-iGluSnFR protein at the attachment site between the cells was examined using HEK293T cells. Plasmids encoding TagBFP-P2A-Pre-Sphere-SF-iGluSnFR or iRFP-P2A-Post-Sphere-SF-iGluSnFR were transfected by electroporation using a Neon Transfection System (Thermo Fisher Scientific). The transfected cells were then mixed, plated on collagen-coated glass bottom dishes, and cultured in DMEM with 10% FBS, 50 U/mL penicillin, and 50 μg/mL streptomycin. 1–2 days after plating, the cells were rinsed with HBSS, and fluorescence was observed using the FluoView FV1000 microscope. TagBFP was excited at a wavelength of 405 nm from a laser diode, iRFP was excited at 635 nm from a laser diode, and Sphere-SF-iGluSnFR was excited at 488 nm from an Ar laser through a DM405/488/559/635 dichroic mirror (Olympus). The fluorescence wavelengths were separated with dichroic mirrors SDM490 and SDM560, and were observed using photomultipliers at wavelengths of 425–475 nm for TagBFP and 500–560 nm for Sphere-SF-iGluSnFR through spectroscopy unit, and 655–755 nm for iRFP through a band pass filter.

### *C. elegans* strains

Worms were cultured at 20°C on nematode growth medium agar plates with *Escherichia coli* OP50 bacteria under standard conditions^41^. Hermaphrodites were used for all experiments.

Codon-optimized Pre-Sphere-SF-iGluSnFR and Post-Sphere-SF-iGluSnFR constructs for expression in *C. elegans* were produced from the codon-optimized iGluSnFR, which was gifted by Dr Loren Looger. Expression markers TagBFP or mCherry were attached to Pre- or Post-Sphere-SF-iGluSnFR, respectively, via an SL2 trans-splicing sequence. The resulting TagBFP-SL2-Pre-Sphere-SF-iGluSnFR and mCherry-SL2-Post-Sphere-SF-iGluSnFR sequences were cloned into Gateway Destination vectors (Thermo Fisher Scientific). Gateway Entry vectors (Thermo Fisher Scientific) with cell-specific promoters were obtained from the Comprehensive Brain Science Network. Plasmids for the AWC neuron-specific expression of TagBFP-SL2-Pre-Sphere-SF-iGluSnFR and for the AIY neuron-specific expression of mCherry-SL2-Post-Sphere-SF-iGluSnFR in *C. elegans* were generated from these vectors using Gateway Cloning Technology (Thermo Fisher Scientific). The transgenic strain was created via microinjection of both vectors into the N2 Bristol strain (wild type).

### Confocal imaging for *C. elegans*

Confocal images of Sphere expression (Fig. 3A) were acquired using the Olympus FluoView FV1000 confocal laser scanning microscopy system with the sample mounted on an inverted microscope (IX81, Olympus) using a 60× oil-immersion objective (UPlanSApo, NA: 1.35, Olympus). Worms were immobilized with 20 mM sodium azide and mounted in a 1% low-melting-point agarose gel (UltraPure, Invitrogen). The TagBFP, mCherry, and Sphere-SF-iGluSnFR proteins were sequentially excited at 405 nm, 559 nm, and 488 nm, respectively, through a DM405/488/559 dichroic mirror (Olympus). The fluorescence was separated with dichroic mirrors, SDM490 and SDM560, and observed using photomultipliers at wavelengths of 425–475 nm for TagBFP, 500–545 nm for Sphere-SF-iGluSnFR, and 575-675 nm for mCherry. Z-stack images were acquired every 1.5 μm, and Z-projection images were created from all of the Z-stack images.

### Glutamate imaging in *C. elegans*

For glutamate imaging, olfactory chips^28^ were used. The worms were stimulated with S-basal buffer and IAA diluted with S-basal buffer (9.2 × 10^−4^ M). Fluorescence imaging was performed using an inverted microscope (IX71, Olympus) equipped with a 20× objective (UCPLFLN 20X, N. A.: 0.7, Olympus) in addition to a 1.6× zoom lens, U-MWIB3 cube (Olympus), an LED light source (SOLA, Lumencor, Beaverton, OR, USA), and a 3CCD camera (C7800-20, Hamamatsu Photonics, Hamamatsu, Japan). Images were acquired every 200 ms, with the same exposure time using AQUACOSMOS software (Hamamatsu Photonics). To reduce noise caused by worm movement, S-basal buffer containing the cholinergic agonist levamisole (2 mM) was used.

The obtained fluorescence intensity data were normalized by the average intensities for the first 2 s, and the time-courses are presented as ΔF/F_0_.

### Adeno-associated virus (AVV) production

The pAAV-hSyn-EGFP vector was a gift from Dr Bryan Roth (Addgene plasmid #50465). TagBFP-P2A-Pre-Sphere-SF-iGluSnFR and mCherry-P2A-Post-Sphere-SF-iGluSnFR sequences were inserted into the pAAV-hSyn-EGFP Vector by replacing the EGFP using Infusion (Takara Bio). AAV serotype 2 was amplified in HEK293T cells by transfecting pAAV.hSynapsin.TagBFP-P2A-Pre-Sphere-SF-iGluSnFR or pAAV.hSynapsin.mCherry-P2A-Post-Sphere-SF-iGluSnFR via the pRC2-mi342 and pHelper vectors, respectively (Takara Bio). The transfected cells were incubated in five 225 cm^2^ flasks for three days. The cells were collected and centrifuged at 1,750 × g for 10 min at 4°C. AAV vectors were subsequently extracted from the pellet using an AAVpro purification Kit (Takara bio). The eluted solution was concentrated using an Amicon Ultra-15 centrifugal filter unit (Takara bio). The titre was measured via real-time PCR using an AAVpro Titration Kit for Real Time PCR Ver. 2 (Takara Bio).

### *In vivo* mouse experiments

Male mice (C57BL/6, 2–3 months old, *N* = 2) were used for the *in vivo* imaging experiments. The skull above the sensorimotor cortex of the left hemisphere was removed (approximately 4 mm in diameter) using a dental drill while under isoflurane (2%–3%) anaesthesia. The diluted viral vectors (500 nL) expressing the TagBFP-P2A-Pre-Sphere-SF-iGluSnFR and mCherry-P2A-Post-Sphere-SF-iGluSnFR constructs were injected at a depth of 300 μm from the cortical surface using a glass pipette. A total of six injections (three for Pre-Sphere-SF-iGluSnFR and three for Post-Sphere-SF-iGluSnFR) were performed on each animal. The surface of the exposed brain was then covered with a cover glass, and secured with dental resin. After ceasing anaesthesia treatment, the animals were allowed to recover in a normal home cage with *ad libitum* access to food and water.

T Three weeks after viral injection, the animals were once again anaesthetized with isoflurane (3%–5% for induction and 1%–3% for imaging experiments), and live imaging experiments using a two-photon microscope (SP8MP, Leica Microsystems, Wetzlar, Germany) equipped with a Ti: sapphire laser (MaiTaiHP, Spectra Physics, Milpitas, CA, USA) were performed to measure fluorescence signals under awake or anesthetized conditions. Two detectors (525/50 nm and 610/75 nm) were used to simultaneously receive the emission signals through a band separator (560/10 nm). To confirm the sensor and marker expression, the excitation wavelength was changed to 850 nm for TagBFP, 910 nm for Sphere-SF-iGluSnFR, and 1020 nm for mCherry at each imaging location. Time-lapse images of Sphere-SF-iGluSnFR fluorescence (1024 × 1024 pixels, 0.4 μm/pixel) were captured from 100 to 300 frames through an objective lens (HCX IRAPOL 25×, NA = 1.00, Leica Microsystems) at a rate of 2.5 s per frame (i.e., 400 Hz scan speed) in the upper layer of cortical layer II/III.

### Analysis of synaptic glutamate transmission

To analyse the *in vivo* time-lapse images, ROIs were selected from blinking fluorescent spots found under awake conditions using the Aquacosmos software (Hamamatsu Photonics). Fluorescent spots that did not appear to be blinking were not selected for analysis. Slight movements of the fluorescent spots, likely due to movements of the animal, were manually corrected by changing the position of the ROIs throughout the time-lapse. Time-courses of the averaged fluorescence intensities at each ROI were calculated and further processed using custom-designed MATLAB programs. The fluorescence intensity time-courses at each ROI were normalized using the most frequent intensity value in the time-course, and the temporal differentiation of the normalized fluorescence intensity (dF/dt) was calculated for each ROI as shown in Fig. 4D. The responses detected from the time-course were plotted as dF/dt. The mean and the standard deviation (SD) of dF/dt were calculated and the values between the means ± (3 × SD) were defined as Noise for each ROI. The mean and SD of the Noise were calculated, and the responses were defined for each ROI as follows:

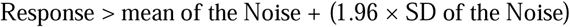

To detect slow responses, the same process was adopted for the time-course processed using a two-frame moving average, and the detected points were also defined as responses, with the exception of the already defined points and the ± 1 frame adjacent to them. When the response was continuous for two or more frames, only the first frame was defined as the response. Transient loss of the fluorescing spots was likely due to animal movement, and was observed at some time points. To exclude the responses detecting fluorescence reappearance after disappearance, data points where the value one frame before was lower than the mean of the Noise – (1.96 × SD of the Noise) were excluded. The detected responses were shown in raster plots (Fig. 4E). The number of responding ROIs was calculated for each frame, and synchronous activity was considered to occur when more than 10 ROIs responded at the same point.

The response of each ROI was clustered using TSclust, an R package for time series clustering^42^. The time-course of dF/dt for all ROIs were inputted and clustered based on the correlations among them.

Correlations between the time-course of normalized fluorescence intensity for each ROI pair were calculated using Pearson correlation coefficients and applying a Fisher Z-transformation.

## Data availability

All data from this study are available upon request.

## Acknowledgement

We want to thank Mr. Yuki Yano of the Faculty of Informatics and Engineering at the University of Electro-Communications for his support with *in vivo* imaging of the mouse brains.

## Author contributions

K.A. designed the study. Y.S. designed and engineered the sensor, and performed fluorescence imaging of the cultured cells. K.A. performed fluorescence imaging in *C. elegans*. H.T., M.T., and M.H. performed the virus injection into the mouse brains. K.M. performed *in vivo* imaging in the mouse brains. R.I. wrote the MATLAB script and analysed *in vivo* imaging data. Y.S. created the figures. Y.S., K.A., and K.M. wrote the manuscript. All authors reviewed and edited the manuscript. K.H. and K.O. supervised all of the work.

## References

1. Denk, W. & Horstmann, H. Serial Block-Face Scanning Electron Microscopy to Reconstruct Three-Dimensional Tissue Nanostructure. PLoS Biol. 2, e329 (2004).

2. Nava Gonzales, C. et al. Systematic morphological and morphometric analysis of identified olfactory receptor neurons in Drosophila melanogaster. eLife 10, e69896 (2021).

3. Dorkenwald, S. et al. Automated synaptic connectivity inference for volume electron microscopy. Nat. Methods 14, 435–442 (2017).

4. White, J. G., Southgate, E., Thomson, J. N. & Brenner, S. The structure of the nervous system of the nematode Caenorhabditis elegans. Philos. Trans. R. Soc. Lond. B. Biol. Sci. 314, 1–340 (1986).

5. Mann, K., Gallen, C. L. & Clandinin, T. R. Whole-Brain Calcium Imaging Reveals an Intrinsic Functional Network in Drosophila. Curr. Biol. 27, 2389-2396.e4 (2017).

6. Kato, S. et al. Global Brain Dynamics Embed the Motor Command Sequence of Caenorhabditis elegans. Cell 163, 656–669 (2015).

7. Ahrens, M. B. et al. Brain-wide neuronal dynamics during motor adaptation in zebrafish. Nature 485, 471–477 (2012).

8. Venkatachalam, V. et al. Pan-neuronal imaging in roaming Caenorhabditis elegans. Proc. Natl. Acad. Sci. U. S. A. 113, E1082–E1088 (2016).

9. Nguyen, J. P. et al. Whole-brain calcium imaging with cellular resolution in freely behaving Caenorhabditis elegans. Proc. Natl. Acad. Sci. 113, E1074–E1081 (2016).

10. Lee, W.-C. A. et al. Anatomy and function of an excitatory network in the visual cortex. Nature 532, 370–374 (2016).

11. Vishwanathan, A. et al. Electron Microscopic Reconstruction of Functionally Identified Cells in a Neural Integrator. Curr. Biol. 27, 2137-2147.e3 (2017).

12. Marvin, J. S. et al. An optimized fluorescent probe for visualizing glutamate neurotransmission. Nat. Methods 10, 162–170 (2013).

13. Wan, J. et al. A genetically encoded sensor for measuring serotonin dynamics. Nat. Neurosci. 24, 746–752 (2021).

14. Sun, F. et al. Next-generation GRAB sensors for monitoring dopaminergic activity in vivo. Nat. Methods 17, 1156–1166 (2020).

15. Patriarchi, T. et al. Ultrafast neuronal imaging of dopamine dynamics with designed genetically encoded sensors. Science 360, eaat4422 (2018).

16. Marvin, J. S. et al. A genetically encoded fluorescent sensor for in vivo imaging of GABA. Nat. Methods 16, 763–770 (2019).

17. Xie, Y. et al. Resolution of High-Frequency Mesoscale Intracortical Maps Using the Genetically Encoded Glutamate Sensor iGluSnFR. J. Neurosci. 36, 1261–1272 (2016).

18. Kim, J. et al. mGRASP enables mapping mammalian synaptic connectivity with light microscopy. Nat. Methods 9, 96–102 (2012).

19. Feinberg, E. H. et al. GFP Reconstitution Across Synaptic Partners (GRASP) Defines Cell Contacts and Synapses in Living Nervous Systems. Neuron 57, 353–363 (2008).

20. Choi, J.-H. et al. Interregional synaptic maps among engram cells underlie memory formation. Science 360, 430–435 (2018).

21. Marvin, J. S. et al. Stability, affinity, and chromatic variants of the glutamate sensor iGluSnFR. Nat. Methods 15, 936–939 (2018).

22. Cabantous, S. et al. A New Protein-Protein Interaction Sensor Based on Tripartite Split-GFP Association. Sci. Rep. 3, 2854 (2013).

23. Pedelacq, J.-D. & Cabantous, S. Development and Applications of Superfolder and Split Fluorescent Protein Detection Systems in Biology. Int. J. Mol. Sci. 20, 3479 (2019).

24. Pisabarro, M. T. & Serrano, L. Rational Design of Specific High-Affinity Peptide Ligands for the Abl-SH3 Domain. Biochemistry 35, 10634–10640 (1996).

25. Wu, J. et al. Genetically Encoded Glutamate Indicators with Altered Color and Topology. ACS Chem. Biol. 13, 1832–1837 (2018).

26. Ashida, K., Hotta, K. & Oka, K. The Input-Output Relationship of AIY Interneurons in Caenorhabditis elegans in Noisy Environment. iScience 19, 191–203 (2019).

27. Ashida, K., Shidara, H., Hotta, K. & Oka, K. Optical Dissection of Synaptic Plasticity for Early Adaptation in Caenorhabditis elegans. Neuroscience 428, 112–121 (2020).

28. Chronis, N., Zimmer, M. & Bargmann, C. I. Microfluidics for in vivo imaging of neuronal and behavioral activity in Caenorhabditis elegans. Nat. Methods 4, 727–731 (2007).

29. Masamoto, K., Fukuda, M., Vazquez, A. & Kim, S.-G. Dose-dependent effect of isoflurane on neurovascular coupling in rat cerebral cortex. Eur. J. Neurosci. 30, 242–250 (2009).

30. Masamoto, K., Kim, T., Fukuda, M., Wang, P. & Kim, S.-G. Relationship between Neural, Vascular, and BOLD Signals in Isoflurane-Anesthetized Rat Somatosensory Cortex. Cereb. Cortex 17, 942–950 (2007).

31. Gui, S. et al. Revealing the Cortical Glutamatergic Neural Activity During Burst Suppression by Simultaneous wide Field Calcium Imaging and Electroencephalography in Mice. Neuroscience 469, 110–124 (2021).

32. Liu, X. et al. Optogenetic stimulation of a hippocampal engram activates fear memory recall. Nature 484, 381–385 (2012).

33. Lobas, M. A. et al. A genetically encoded single-wavelength sensor for imaging cytosolic and cell surface ATP. Nat. Commun. 10, 711 (2019).

34. Keller, J. P. et al. In vivo glucose imaging in multiple model organisms with an engineered single-wavelength sensor. Cell Rep. 35, 109284 (2021).

35. Nasu, Y. et al. A genetically encoded fluorescent biosensor for extracellular l-lactate. Nat. Commun. 12, 7058 (2021).

36. Dong, A. et al. A fluorescent sensor for spatiotemporally resolved imaging of endocannabinoid dynamics in vivo. Nat. Biotechnol. 1–12 (2021).

37. Chen, T.-W. et al. Ultrasensitive fluorescent proteins for imaging neuronal activity. Nature 499, 295–300 (2013).

38. Dana, H. et al. High-performance calcium sensors for imaging activity in neuronal populations and microcompartments. Nat. Methods 16, 649–657 (2019).

39. Guertin, P. A. The mammalian central pattern generator for locomotion. Brain Res. Rev. 62, 45–56 (2009).

## References

40. Percie du Sert, N. et al. The ARRIVE guidelines 2.0: Updated guidelines for reporting animal research*. J. Cereb. Blood Flow Metab. 40, 1769–1777 (2020).

41. Brenner, S. The genetics of Caenorhabditis elegans. Genetics 77, 71–94 (1974).

42. Montero, P. & Vilar, J. A. TSclust□: An R Package for Time Series Clustering. J. Stat. Softw. 62, 1–43 (2014).

