## Supplementary Figure for "Genetically encoded sensors for analysing neurotransmission among synaptically-connected neurons"

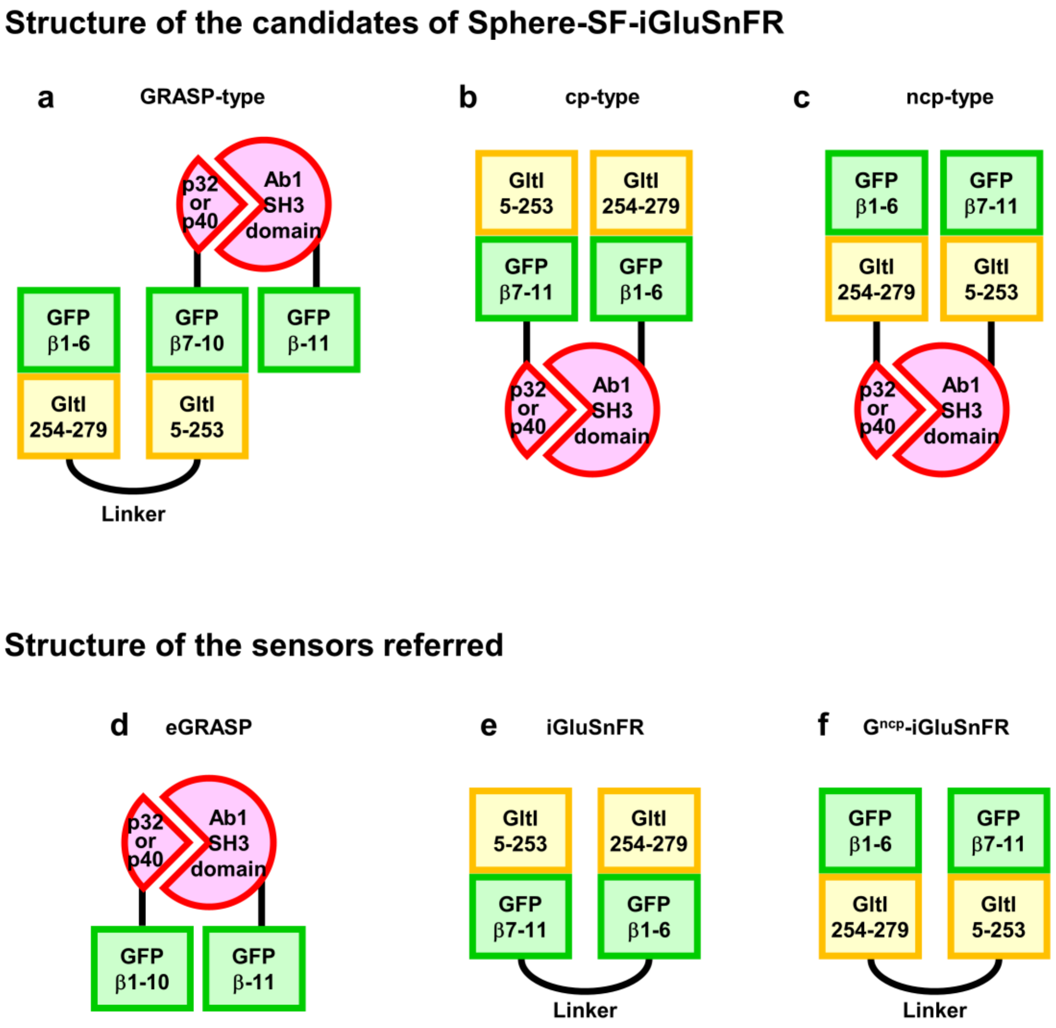


Supplementary Figure 1. Schematic structure of SF-iGluSnFR, $G^{ncp}$-iGluSnFR, eGRASP, and Sphere-SF-iGluSnFR candidates. Green boxes: GFP fragments; yellow boxes: fragments of the glutamate sensor domain; and red sectors: selective binding domains.

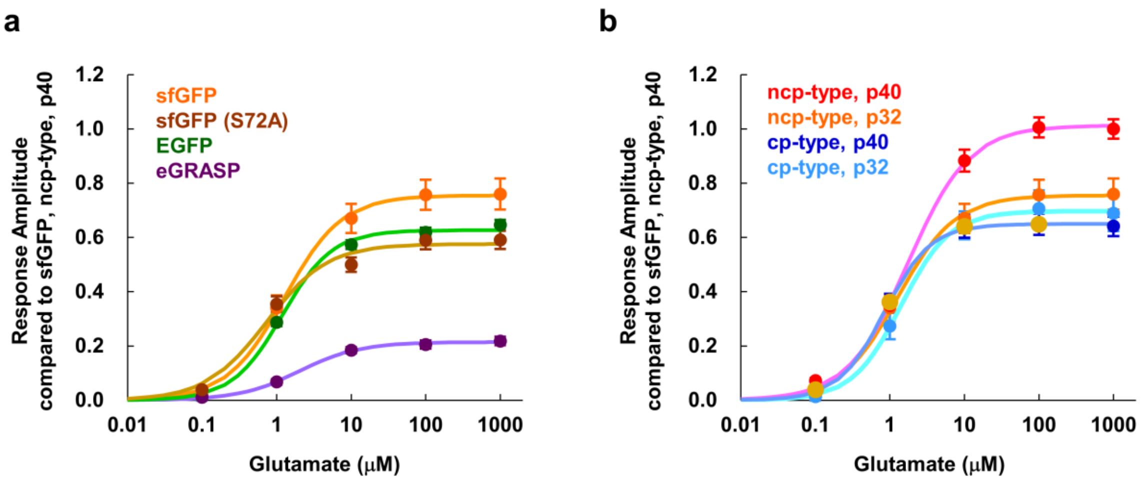


Supplementary Figure 2. Comparison of GFP variants and structures in the Sphere-SF-iGluSnFR system. GFP variants used in the sensor, the composition of the sensor (circularly permutated [cp-type] or non-circularly permutated [ncp-type]), and variants of the selective binding domains targeted to the AB1 SH3 domain (p32 or p40) were examined. The responses were measured by expressing both fragments of the Sphere-SF-iGluSnFR protein in HeLa cells. Observed response amplitudes were normalized using the saturation amplitude of Sphere-SF-iGluSnFR, all with sfGFP, ncp-type, and p40; red in (b). (a) Comparison of sensor responses with different GFP variants (ncp-type with a p32 domain). Orange: sfGFP (*n* = 57 cells from 6 different experiments); brown: sfGFP with a S72A mutation (*n* = 73 cells from 6 different experiments); green: EGFP (*n* = 46 cells from 5 different experiments); and purple: GFP used in eGRASP (*n* = 46 cells from 5 different experiments). (b) Comparison of the sensor types (shown in Supplementary Fig. S1) and binding domains (all sfGFP). Red: ncp-type with p40 (*n* = 104 cells from 8 different experiments); orange: ncp-type with p32 (*n* = 57 cells from 6 different experiments); blue: cp-type with p40 (*n* = 67 cells from 7 different experiments); and light blue: cp-type with p32 (*n* = 51 cells from 8 different experiments). Error bars: SEM.

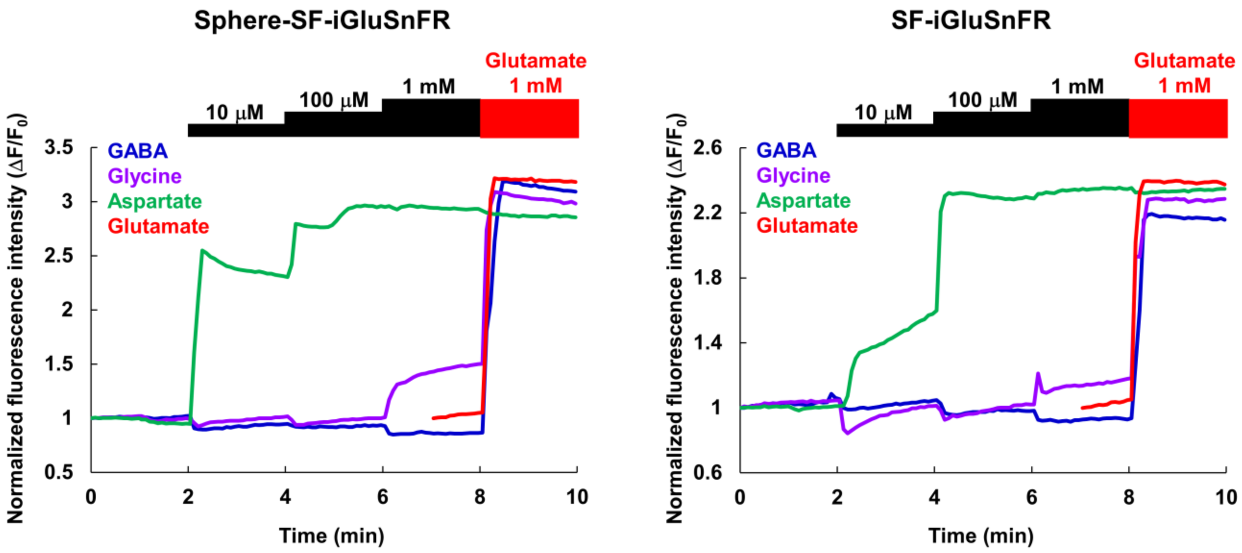


Supplementary Figure 3. Comparison of the ligand selectivity of the Sphere-SF-iGluSnFR (left) compared to the original SF-iGluSnFR (right). The averaged time-course of the sensor responses to the stepwise application of GABA (blue; $n$ = 71 cells from 5 different experiments for Sphere-SF-iGluSnFR, and $n$ = 58 cells from 3 different experiments for the original SF-iGluSnFR), glycine (purple; $n$ = 71 cells from 5 different experiments for Sphere-SF-iGluSnFR and $n$ = 56 cells from 3 different experiments for the original SF-iGluSnFR), and aspartate (green; $n$ = 69 cells from 5 different experiments for Sphere-SF-iGluSnFR and $n$ = 56 cells from 3 different experiments for the original SF-iGluSnFR) followed by 1 mM glutamate were compared to the responses to only glutamate (red; $n$ = 78 cells from 5 different experiments for Sphere-SF-iGluSnFR and $n$ = 55 cells from 3 different experiments for the original SF-iGluSnFR). These two sensors showed similar responses.

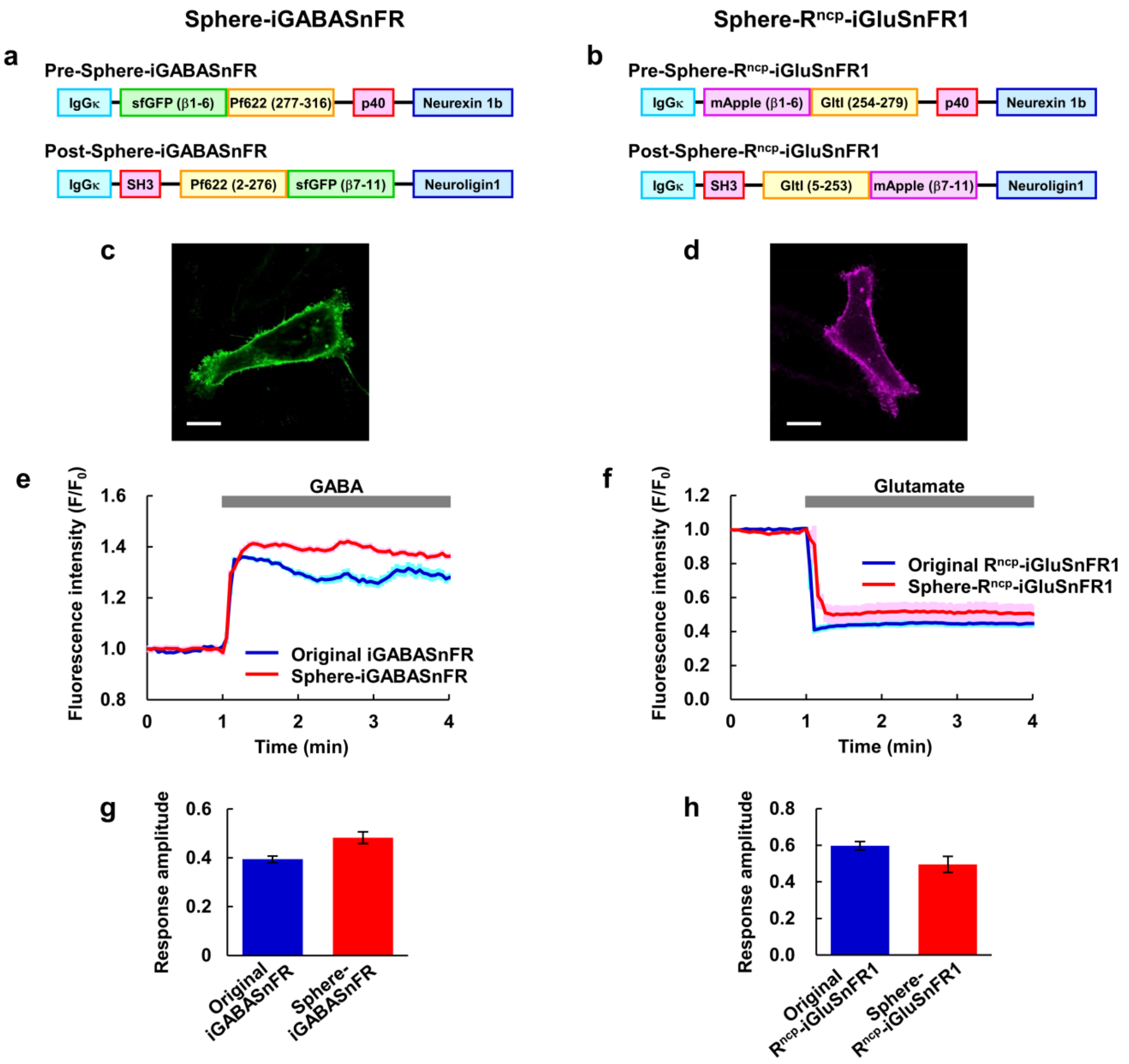


Supplementary Figure 4. The Split Protein HEmispheres for REconstitution (Sphere) technique can be applied to iGABASnFR and $R^{ncp}$-iGluSnFR sensors. (a, b) Schematic of Pre- and Post-Sphere-iGABASnFR and Pre- and Post-Sphere-$R^{ncp}$-iGluSnFR1 constructs. (c, d) Fluorescence images of reconstituted Sphere-iGABASnFR and Sphere-$R^{ncp}$-iGluSnFR1 expressed on HeLa cells. Scale bar: 20 μm. (e) Time-course of the normalized fluorescence of the original iGABASnFR (blue: $n$ = 14 cells from 3 different experiments) and Sphere-iGABASnFR (red: $n$ = 14 cells from 4 different experiments) in response to GABA (1 mM). (f) Time-courses of the normalized fluorescence of the original $R^{ncp}$-iGluSnFR (blue: $n$ = 11 cells from 4 different experiments) and Sphere-$R^{ncp}$-iGluSnFR (red: $n$ = 6 cells from 3 different experiments) in response to glutamate (1 mM). (g) Comparison of response amplitudes of the original sensor and Sphere-iGABASnFR. (h) Comparison of the response amplitudes of the original sensor and Sphere-$R^{ncp}$-iGluSnFR1. The response amplitudes in (g) and (h) were calculated as the difference between the average normalized fluorescence within ten frames immediately before the application of GABA or glutamate, and the maximum or minimum normalized fluorescence in 1–4 min. Error bars in (e)–(h): SEM.

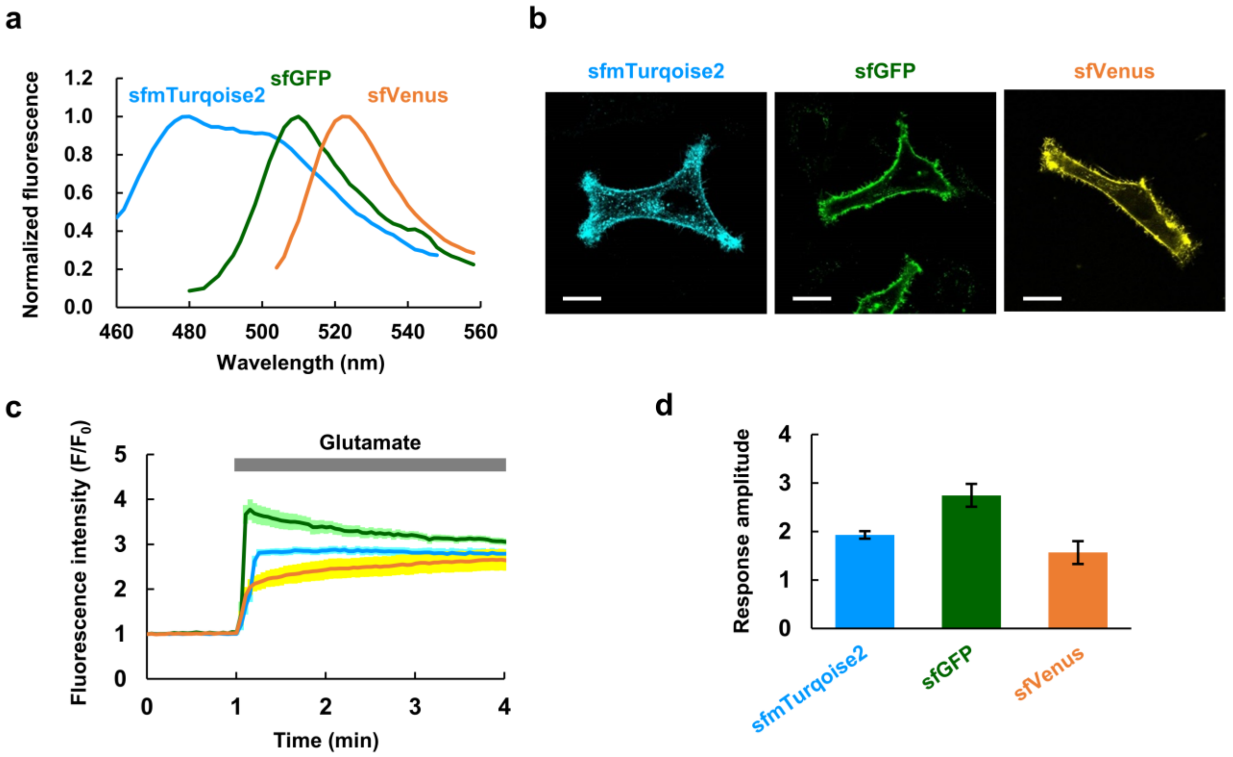


Supplementary Figure 5. Colour variants of the Sphere-SF-iGluSnFR construct. The mutations changing sfGFP to sfmTurquoise2 were created at T65S, Y66W, and S72A (GFP numbering). Mutations in the linker between sfmTurquoise2 and GltI were also put according to sfmTurquoise20iGluSnFR. The mutations changing sfGFP to sfVenus were created at F64L, T65G, S72A, and T203Y (GFP numbering). The normalized fluorescence spectrum (a) and fluorescence images (b) were measured in HeLa cells. Scale bar: 20 µm. (c) Time-courses of normalized fluorescence intensity as a result of Sphere-sfmTurquoise2-iGluSnFR (cyan; $n$ = 23 cells from 3 different experiments), Sphere-SF-iGluSnFR (green; $n$ = 13 cells from 4 different experiments), and Sphere-sfVenus-iGluSnFR (orange; $n$ = 11 cells from 4 different experiments) in response to glutamate (1 mM). (d) A comparison of response amplitudes. Error bars in (c) and (d): SEM.

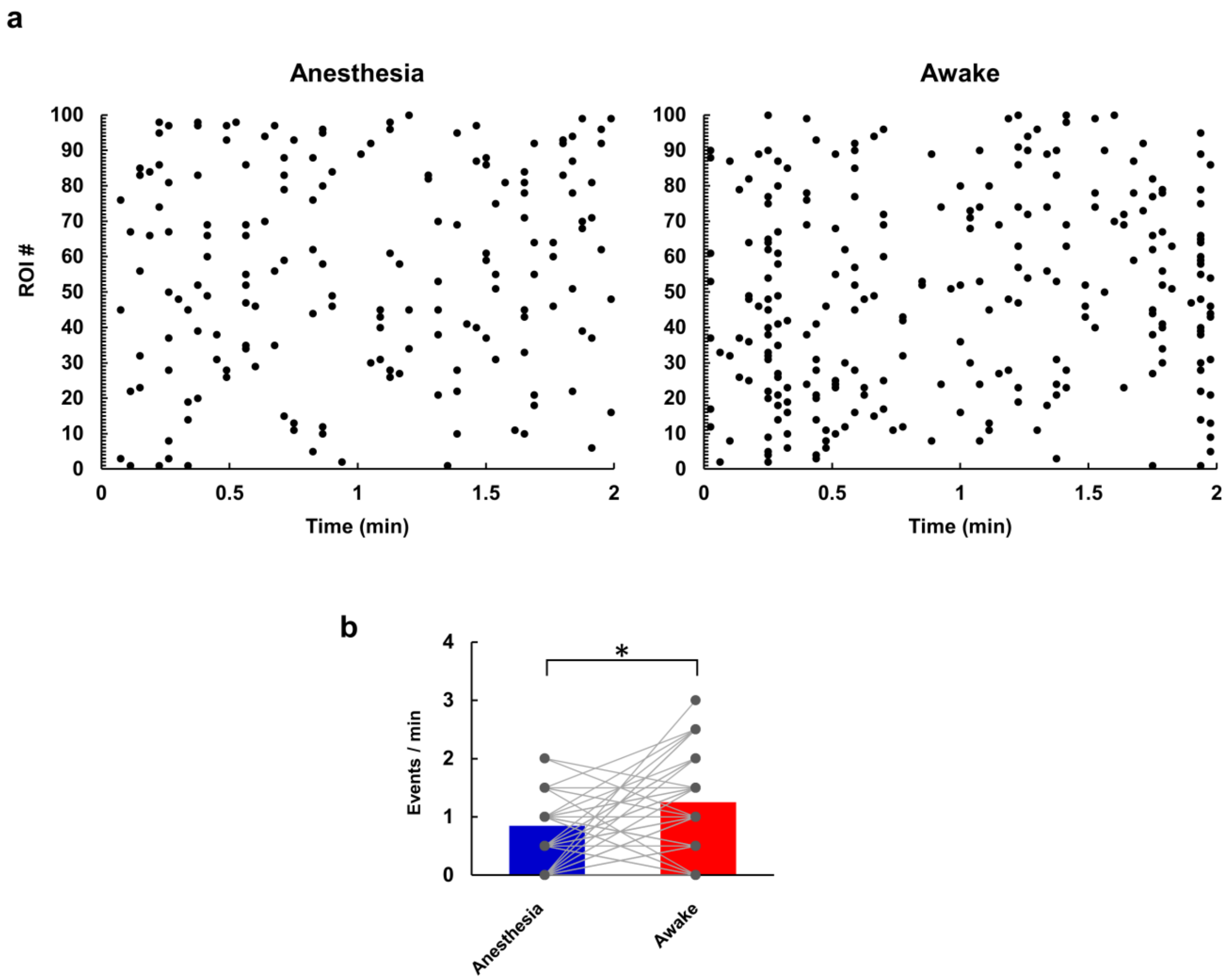


Supplementary Figure 6. The *in vivo* imaging of Sphere-SF-iGluSnFR in another mouse. (a) Raster plots of spontaneous synaptic activities under anaesthetised and subsequent awake conditions (100 ROIs from a single time series). (b) Comparison of the response frequencies of each region of interest (ROI; grey dots) and the overall average (coloured bars). A significantly higher response frequency was observed while the mouse was under awake conditions compared to anaesthetised conditions. * indicates $P < 0.05$, Student's $t$-test.
